# Carbohydrate consumption drives adaptive mutations in *Escherichia coli* associated with increased risk for systemic infection

**DOI:** 10.1101/2025.03.26.645536

**Authors:** Aaron L. Hecht, Nadim Mahmud, Subhan Chaudhry, Jacob Y. Cao, Grace Purvis Branigan, Junhee Lee, Erin Theiller, Manuela Roggiani, Elliot S. Friedman, Lindsay Herman, Bianca E. Galis, Steven M. Jones, Paul J. Planet, Joseph P. Zackular, David E. Kaplan, Marina Serper, K. Rajender Reddy, Ahmed M. Moustafa, Mark Goulian, Gary D. Wu

**Affiliations:** Division of Gastroenterology and Hepatology, Perelman School of Medicine, University of Pennsylvania, Philadelphia, PA; Gastroenterology Section, Corporal Michael J. Crescenz VA Medical Center, Philadelphia PA; Hackensack Meridian School of Medicine, Nutley, NJ, USA; Department of Medicine, University of Cambridge School of Clinical Medicine, Cambridge Biomedical Campus, Cambridge, UK; Center for Microbial Medicine, Children’s Hospital of Philadelphia; Philadelphia, PA, USA; Division of Gastroenterology, Hepatology, and Nutrition, The Children’s Hospital of Philadelphia, Philadelphia, PA, USA; Department of Biology, University of Pennsylvania, Philadelphia, PA, USA; Department of Pediatrics, Perelman School of Medicine, University of Pennsylvania, Philadelphia, PA, USA; Division of Pediatric Infectious Diseases, Children’s Hospital of Philadelphia, Philadelphia, PA, USA; Department of Pathology and Laboratory Medicine, Perelman School of Medicine, University of Pennsylvania; Philadelphia, PA, USA; Department of Pathology and Laboratory Medicine, Children’s Hospital of Philadelphia; Philadelphia, PA, USA

## Abstract

Dissemination of organisms from the gut microbiota is a major contributor to sepsis and critical illness. Patients with cirrhosis are prone to systemic infections and are commonly prescribed the carbohydrate lactulose to manage hepatic encephalopathy (HE) ^1^. Commensal metabolism of lactulose is believed to reduce pathobiont colonization through short-chain fatty acid production, but its direct effects on gut pathobionts remain unexplored ^2^. Here, we show that lactulose consumption unexpectedly selects for mutations in *Escherichia coli lactose (lac)* operon regulation, enhancing its metabolic fitness and colonization capacity. This is mediated by selection for constitutive expression of the *lac* operon through mutations in its regulatory components. Using *in vitro* systems, murine models, and clinical samples, we demonstrate that these mutations enable *E. coli* to exploit lactulose as a carbon source, bypassing host carbohydrate metabolism and increasing its intestinal colonization. Despite its long-standing use in HE treatment, we find that lactulose has a paradoxical association with risk of infection hospitalization in patients with cirrhosis in a large epidemiologic study. The emergence of lactulose-adapted *E. coli* strains could be suppressed by a dietary oligosaccharide that competitively inhibits lactulose uptake. These findings reveal a mechanism by which dietary substrates exert selective pressure on the microbiome, with implications for diet-based strategies to modulate microbial evolution and infection risk.

## Introduction

Systemic infections originating from the gut microbiota are a significant cause of morbidity and mortality in susceptible individuals. Patients with liver cirrhosis, where infection-related mortality approaches 50%, are particularly vulnerable to spontaneous bacterial peritonitis (SBP) and bacteremia due to enteric bacterial translocation ^1^. Emerging evidence highlights the role of microbiota composition in influencing SBP risk, with reduced diversity identified as a major contributing factor ^2^. Progressive liver disease is associated with decreased microbiota diversity and the proliferation of *Enterobacteriaceae*, including the pathobiont *Escherichia coli*, a dysbiotic state that predisposes to infection ^3–8^. Notably, *E. coli* is among the most common bacterial isolates in bloodstream infections and SBP in this population, underscoring the link between increased *E. coli* colonization in dysbiosis and disseminated infection ^9–11^. Despite this, the environmental and bacterial factors driving disseminated infections remain poorly understood.

Dietary modifications have demonstrated clinical efficacy in mitigating dysbiosis, with extensive research examining the effects of diet on bacterial commensals ^12–21^. However, the impact of diet on *Enterobacteriaceae* remains underexplored. Recent work suggests that the colon is a carbohydrate-limited environment for *Enterobacteriaceae* due to nutrient absorption by the small intestine of the mammalian host ^22^. Lactulose, an artificial disaccharide that bypasses host absorption, has been the standard of care therapy for over 50 years as a treatment for hepatic encephalopathy (HE), a common complication of cirrhosis ^23,24^. The delivery of lactulose to the colonic microbiota promotes the growth of commensal bacterial genera such as *Lactobacillus* and *Bifidobacterium*, leading to the increased production of short-chain fatty acids (SCFAs) ^25–27^. Little attention has been devoted to the impact of lactulose directly on *Enterobacteriaceae* and in the context of dysbiosis.

Our previous findings revealed that lactulose use in cirrhotic patients is associated with an increased abundance of *E. coli* in the gut microbiota ^28^. As *E. coli* has not been reported to robustly metabolize lactulose, we sought to elucidate the mechanism underlying this association. Here, we demonstrate that lactulose enhances *E. coli* colonization in the gut by selecting for mutants that constitutively express the lactose (*lac*) operon. Using *in vitro* cultures, mouse models, and human bacterial isolates, we show that lactulose exerts a strong and rapid selective pressure, driving genetic adaptation of *E. coli* in the mammalian microbiota. Furthermore, a large cohort study revealed that lactulose is associated with an increased risk of systemic infection in cirrhotic patients specifically with *E. coli*. Notably, the expansion of these lactulose-adapted mutants can be suppressed by competitive inhibition of the *lac* permease through an alternative dietary carbohydrate. These findings demonstrate that dietary carbohydrates can inadvertently drive microbiome evolution, increasing the risk of gut pathobiont colonization and potentially for systemic infection.

## Results

### Lactulose treatment selects for lactulose-metabolizing *Enterobacteriaceae* in the human gut microbiota

A previous study demonstrated a positive correlation between lactulose treatment and *E. coli/Enterobacteriaceae* abundance in a cross-sectional cohort of patients with cirrhosis, called Acute-on-Chronic Liver Failure with Gut Microbiota-Targeted Assessment and Treatment (ACCLIMATE) ^28^. To investigate the mechanistic basis of this association, we isolated *Enterobacteriaceae* strains from stool samples of ACCLIMATE participants (**Fig. 1a**). Metabolic characterization of these isolates revealed a significant positive correlation between patient prescription of lactulose and the prevalence of lactulose-metabolizing *Enterobacteriaceae* (**Fig. 1b; Supplemental Table 1**). Strikingly, none of the *Enterobacteriaceae* strains isolated from the five subjects who had not received lactulose demonstrated the ability to metabolize lactulose.

**Figure 1.**
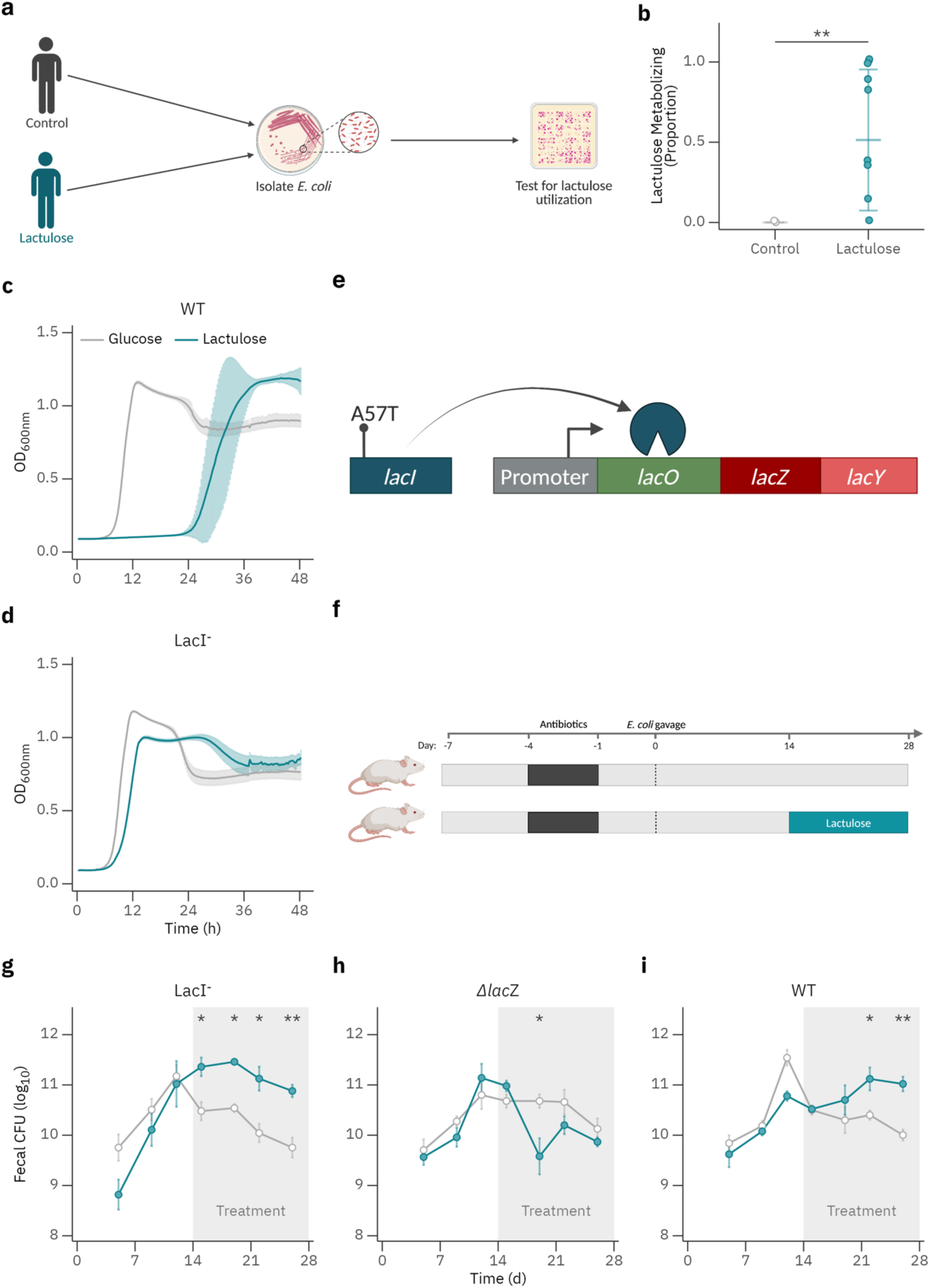
Lactulose consumption selects for lactulose metabolism in gut pathobionts. **A.** Diagram of study design. *Enterobacteriaceae* were isolated from patients with cirrhosis and tested for lactulose metabolism. **B.** Proportion of isolates that were lactulose-metabolizing was quantified and subcategorized by patient treatment with lactulose (n=10-12 isolates per patient; n=11 subjects in lactulose treated, n=5 in lactulose untreated groups). **C and D.** Growth curves of WT (**C**) and LacI^-^ (**D**) *E. coli* strains with glucose or lactulose as a sole carbon source (n=3 wells per group; results are representative of three independent experiments). **E.** Depiction of the *lac* operon with the spontaneous mutation in *lacI* noted. **F.** Diagram of murine study design. Mice were colonized with *E. coli* and treated with or without lactulose in the drinking water. **G-I.** Fecal CFU of mice colonized with LacI^-^ (**G**), WT (**H**), and Δ*lacZ* (**I**) *E. coli* (n=4-5 mice per group; representative of 3 independent experiments). Data shown as mean ± s.d. Statistical analysis was performed using Student’s t-test (**B**), or multiple t-tests with Holm-Sidak correction for multiple comparisons (**G-I**). *p<0.05, **p<0.01.

This finding suggests that the relationship between lactulose consumption and *E. coli* abundance in the gut microbiota is mediated by bacterial adaptation to lactulose metabolism. These results establish a direct link between lactulose treatment and the selection of metabolically specialized *Enterobacteriaceae* within the human gut.

### Mutations in the *lac* operon repressor enable lactulose metabolism in *E. coli*

To investigate how lactulose impacts *E. coli* growth, we tested a previously characterized mouse commensal strain (MP1) in minimal media ^29^. Growth on lactulose exhibited a pronounced lag phase, with exponential growth initiating approximately 24 hours post-inoculation, in contrast to immediate growth on glucose (**Fig. 1c**). This delayed onset suggested the emergence of adaptive mutants. Supporting this hypothesis, subclones isolated from the population demonstrated rapid growth on lactulose in subsequent cultures (**Fig. 1d**). Using MacConkey agar supplemented with lactose or lactulose, we confirmed that the subclones could ferment both sugars, while the wild-type (WT) strain fermented only lactose (**Extended Data Fig. 1a**). Whole genome sequencing (WGS) of the lactulose-adapted subclones identified a missense single nucleotide polymorphism (SNP) in *lacI* (A57T), the gene encoding the lactose operon repressor (**Fig. 1e**). Complementation with WT *lacI* suppressed the mutant phenotype, confirming the role of this mutation in lactulose metabolism (**Extended Data Fig. 1b**). Phage transduction of the mutation into the WT background further validated its causative role (data not shown). These results reveal that mutations in the *lacI* repressor enable *E. coli* to metabolize lactulose effectively, and we refer to these mutants as LacI^-^ clones ^30^.

The *E. coli lac* operon is classically thought to have evolved for lactose metabolism, for which its function has been thoroughly described ^30^. As the *lac* repressor regulates expression of beta-galactosidase (BG) and *lac* permease, encoded by the *lacZ* and *lacY* genes in the *lac* operon respectively, we suspected this metabolic unit was responsible for import and hydrolysis of lactulose. Indeed, deletion of *lacZ* (*ΔlacZ*) abolished growth on lactulose (**Extended Data Fig. 1c**). Competitive inhibition of LacY function with Thiodigalactoside (TDG) delayed growth of the WT strain and LacI^-^ mutant (**Extended Data Fig. 1d, e**). To investigate the mechanism underlying the *lacI* mutation, we assessed BG activity. The LacI^-^ mutants exhibited constitutive BG activity, while the WT strain required induction with isopropyl β-d-1-thiogalactopyranoside (IPTG) to express BG (**Extended Data Fig. 1f**). We confirmed that this effect on BG expression was due to loss of transcriptional repression of *lacZ*, which also extended to *lacY* (**Extended Data Fig. 1g, h**). In summary, lactulose exposure selects for mutations in *lacI*, enabling metabolic adaptation of *E. coli* through deregulation of pre-existing pathways. These findings highlight the evolutionary flexibility of bacterial metabolic regulation in response to an environmental pressure imposed by a carbohydrate.

### A consumed carbohydrate drives selection for metabolic mutants in the gut microbiota

We previously demonstrated that *Enterobacteriaceae* colonization in the gut is limited by a paucity of available carbohydrates in the colon ^22^. This scarcity results from host absorption of simple carbohydrates in the small intestine and the restricted metabolic range of *Enterobacteriaceae* for complex dietary carbohydrates. Lactulose, a non-absorbed disaccharide, bypasses host metabolism and becomes an accessible carbohydrate for the gut microbiota. To test if the LacI^-^ mutation confers enhanced growth in the gut during lactulose exposure, we employed a previously established mouse model of dysbiosis induced by antibiotics and a fiber-free diet ^22^. Mice were colonized with WT, LacI^-^, or Δ*lacZ E. coli* strains with or without lactulose supplementation and fecal colony forming units (CFU) were monitored (**Fig. 1f**). The LacI^-^ strain demonstrated a 10-fold increase in colonization within 24 hours of lactulose treatment compared to controls, a difference that persisted throughout the 2-week exposure (**Fig. 1g**). This effect was lost for the Δ*lacZ* strain, confirming that lactulose response in the gut is dependent upon BG expression (**Fig. 1h**). In contrast, colonization of the WT strain was initially unaffected by lactulose during the first 24 hours but progressively increased over the subsequent 2 weeks of lactulose treatment (**Fig. 1i**). We hypothesized that this delayed increase in WT colonization was driven by the selection of lactulose-metabolizing mutants. Supporting this, 22 of 25 *E. coli* isolates from the stool of WT colonized, lactulose-treated mice displayed enhanced growth on lactulose and constitutive BG expression (**Extended Data Fig. 2a-d**). Characterization and sequencing of the *lacI* gene from representative clones revealed three distinct *lacI* loss-of-function SNPs emerging within a single cage (**Extended Data Fig. 2e and f; Supplemental Table 2**). These findings demonstrate that the availability of a simple carbohydrate like lactulose imposes a strong selective pressure on *E. coli*, enabling the rapid outgrowth of metabolic mutants in the gut microbiota of mice.

### Constitutive *lac* operon expression is counter selected during intestinal colonization

Previous studies have shown that constitutive *lac* operon expression imposes a fitness cost due to dissipation of the proton gradient via Lac permease (LacY) substrate-proton symport ^31^. While our findings demonstrate that lactulose exposure selects for lactulose-metabolizing *E. coli* in the gut microbiota, it remained unclear whether the fitness cost of constitutive LacY expression would lead to a selective disadvantage in the absence of lactulose. During colonization by the LacI^-^ strain, we observed the emergence of lactose-negative colonies in the control groups not exposed to lactulose (**Extended Data Fig. 3a**). Over the 4-week experiment, this lactose-negative population became dominant (**Extended Data Fig. 3b**). Characterization of isolates from these mice revealed a complete loss of BG activity and an inability to metabolize lactulose (**Extended Data Fig. 3c-f**). Sequencing of a representative clone from each mouse revealed identical nonsense mutations in *lacZ* at codon 62 (**Supplemental Table 2**). These results demonstrate that constitutive *lac* operon expression is counter selected in the mammalian gut in the absence of lactulose, potentially due to the associated fitness costs. This finding suggests that the persistence of *lac* operon mutants in *E. coli* depends on continuous exposure to lactulose, emphasizing the dynamic nature of microbial adaptation in response to environmental pressures.

### *Lac* operon regulatory mutation provides a competitive advantage during lactulose exposure in the gut microbiota

Our observations suggest that constitutive *lac* operon expression offers a fitness advantage when a suitable substrate is available but imposes a fitness cost in its absence. To test this, we performed co-colonization experiments using marked strain pairs ^29^. *In vitro* competitions confirmed that the GFP and mCherry markers did not affect growth, consistent with previous studies (**Extended Data Fig. 4a**) ^29^. *In vivo* pairwise competitions revealed a clear selective advantage for the WT and Δ*lacZ* in the absence of lactulose (**Fig. 2a, b; Extended Data Fig. 4b-e**). However, in the presence of lactulose, the LacI^-^ strain exhibited significantly enhanced fitness compared to both WT and Δ*lacZ* strains. Together, these findings underscore the cost-benefit trade-off of constitutive *lac* operon expression and highlight the plasticity of the microbiota in response to consumed substrates.

**Figure 2.**
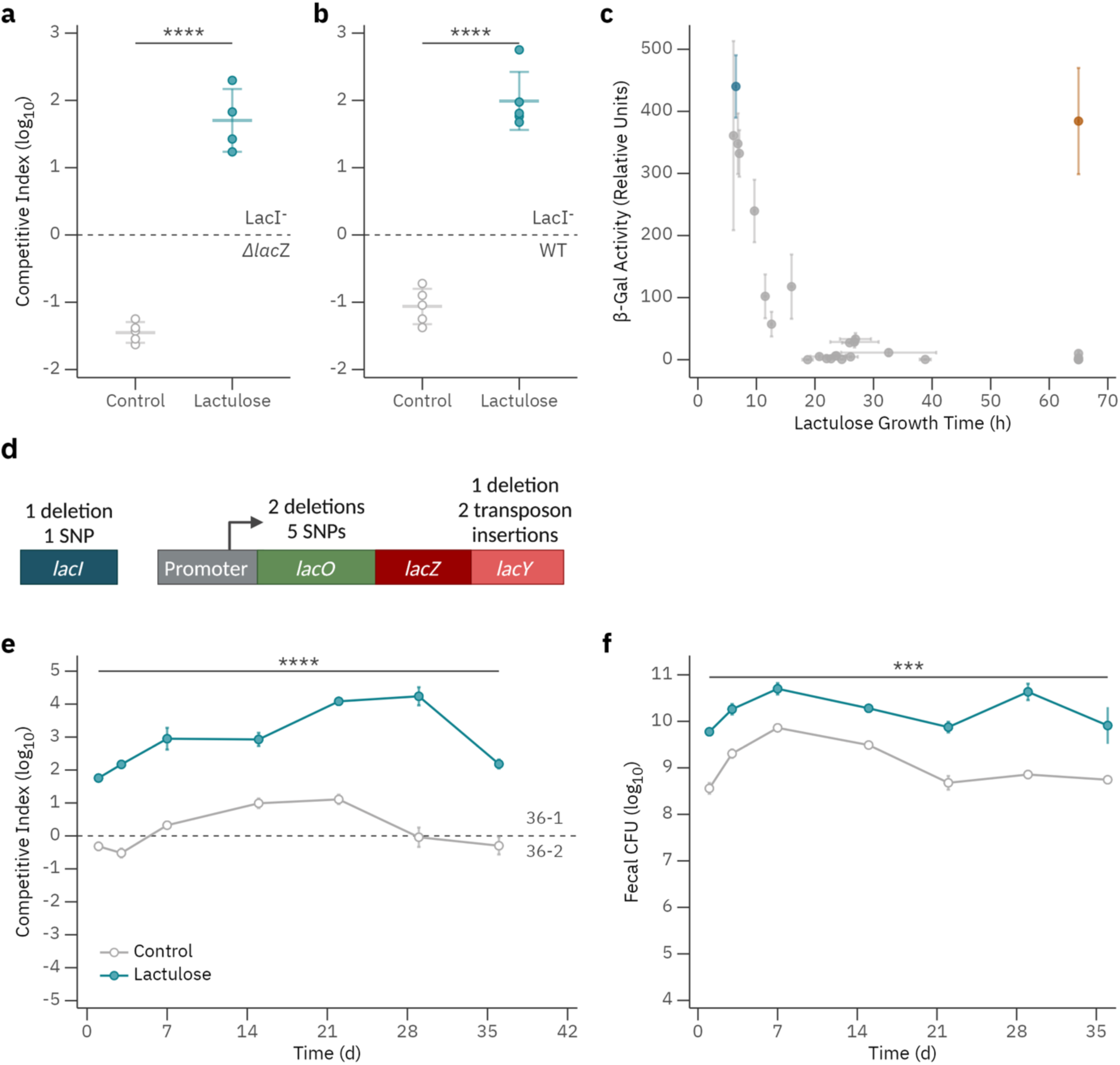
Lactulose treatment is associated with *lac* operon regulatory mutants in the human microbiota. **A and B.** Competitive index between the following strain pairs after 1 week of treatment: Δ*lacZ* and LacI^-^ (**A**); WT and LacI^-^ (**B**), with LacI^-^ displayed as the numerator (n=4-5 mice per group; results representative of two independent experiments). **C.** Correlation between BG activity in the absence of an inducer and growth time on lactulose as a sole carbon source for each phenotypically unique isolate. Outlier strain 36-2 is highlighted in orange and 36-1 in dark blue. (n=6 for BG assay, n=3 for growth time; data representative of two independent experiments). **D.** Summary of *lac* operon and regulatory variants identified from WGS of patient *E. coli* and *K. pneumoniae* isolates. **E.** Competitive index of strains 36-1 and 36-2 after co-colonization of mice with and without lactulose treatment (n=5 mice per group; data representative of two independent experiments). **F**. Fecal CFU of strain 36-1 with and without lactulose treatment (n=5 mice per group; data representative of two independent experiments). Data shown as mean ± s.d. Statistical analysis was performed using Student’s t-test (**A and B**), or multiple t-test with Holm-Sidak correction for multiple comparisons (**E and F**). ***p<0.001, ****p<0.0001.

### Lactulose treatment selects for *lac* operon mutations in cirrhotic patients’ gut microbiota

Based on our findings, we hypothesized that lactulose exerts selective pressure favoring lactulose metabolism by *Enterobacteriaceae* in the human microbiome. To test this, we performed phenotyping and whole genome sequencing (WGS) of unique isolates from the ACCLIMATE study (**Fig. 1a,b**). The majority of isolates were *Escherichia coli* (**Extended Data Fig. 5a**), with a smaller proportion belonging to *Klebsiella* and *Enterobacter* species. Quantification of basal BG activity and growth on lactulose revealed a statistically significant correlation, indicating that altered *lac operon* regulation underlies lactulose metabolism in these strains (**Fig. 2c**). Further analysis of isolates with elevated BG activity identified *E. coli* strains from four separate patients with constitutive *lac* operon expression and rapid lactulose growth (**Extended Data Fig. 5b,c**). WGS revealed mutations in either *lacI* or *lacO*, consistent with previously described disruptions in *lac* operon regulation (**Fig. 2d**; **Supplemental Table 2**) ^32–34^. Similarly, three *Klebsiella pneumoniae* isolates from lactulose-treated patients exhibited elevated BG activity and rapid lactulose metabolism. Although the *K. pneumoniae lac* operon is less well characterized, we identified a shared SNP in *lacO* that may contribute to this phenotype (**Supplemental Table 2**) ^33^. No isolates with elevated BG activity or rapid lactulose growth were found in patients not exposed to lactulose, supporting the role of lactulose as a selective force for these mutants.

To further investigate how selective pressure from lactulose shapes *E. coli* adaptation *in vivo*, we focused on a unique patient case from the ACCLIMATE cohort in which two closely related *E. coli* strains with divergent lactulose-metabolizing phenotypes were isolated. This provided an opportunity to dissect the genetic basis of lactulose metabolism and to study the competitive dynamics of strains with differential metabolic capabilities in response to lactulose exposure. One *E. coli* isolate (36-2) from a lactulose-treated patient displayed elevated basal BG activity but was unable to metabolize lactulose (**Fig. 2c, orange isolate**). This strain represented a minority subpopulation, while the dominant strain (36-1) from the same patient metabolized lactulose and exhibited similarly high basal BG activity (**Supplemental Table 1, Fig 2c, dark blue isolate**). Phylogenetic analysis revealed high relatedness between these strains (**Extended Data Fig. 5a**). Alignment of their shared *lac* operon sequences identified a *lacO* deletion in 36-2, likely driving the elevated basal BG activity due to the loss of the LacI dependent repression of *lacZ* expression (**Extended Data Fig. 5d**). However, 36-2 also carried a frameshift mutation in *lacY*, causing a truncated *Lac* permease, which would prevent lactulose import (**Extended Data Fig. 5e**). Strain 36-1 encodes an additional *lac* operon and *lacI*, but the similarity in basal BG expression between the strains suggests this is not a significant contributor to basal BG expression. Colonization dynamics were explored by marking the strains with distinct antibiotic resistance cassettes. *In vitro* competitions demonstrated a fitness advantage of 36-1 in the presence of lactulose (**Extended Data Fig. 6a**). *In vivo,* competition revealed a significant advantage for 36-1 in the presence of lactulose but similar colonization in its absence (**Fig. 2e**; **Extended Data Fig. 6b,c**). When inoculated independently, 36-1 displayed an anticipated increase in colonization during lactulose treatment (**Fig. 2f**).

Unexpectedly, 36-2 colonization also increased in lactulose-treated mice, accompanied by the emergence of lactulose-positive colonies (**Extended Data Fig. 6d, e**). Sequencing of these colonies revealed two unique insertions in *lacY* that restored the reading frame, enabling lactulose metabolism in separate mice (**Extended Data Fig. 6f**). Co-colonization experiments with 36-1 suppressed these reversion mutants, suggesting competitive inhibition (**Extended Data Fig. 6g**). These findings underscore the role of dietary carbohydrates as strong selective forces in shaping microbiota metabolic capacity.

To test if lactulose selection is common among human-associated *E. coli*, we exposed a library of pediatric bacteremia isolates to lactulose. Initially, 64 out of 90 strains demonstrated growth similar to a reference WT strain (**Extended Data Fig. 7a**). Serial growth in lactulose revealed that most isolates acquired growth rates resembling the LacI^-^ control strain (**Extended Data Fig. 7b**). These data suggest that lactulose exposure readily selects for lactulose metabolism across diverse, disease-associated *E. coli*.

### Lactulose treatment is associated with increased risk of *E. coli* disseminated infections in patients with cirrhosis

Lactulose is one of the most commonly prescribed medications for patients with cirrhosis to manage HE. This population faces high rates of morbidity and mortality due to disseminated infections ^1^. Previous studies have identified *E. coli* as one of the most frequently isolated organisms in both ascites fluid and bloodstream infections in cirrhotic patients ^10,11^. This led us to hypothesize that increased colonization of *E. coli* mutants during lactulose treatment might elevate the risk of bacterial translocation and disseminated infections in these patients. To test this hypothesis, we conducted a retrospective analysis using the Veterans Outcomes and Costs Associated with Liver Disease (VOCAL) cohort ^35,36^, which includes over 100,000 patients with cirrhosis (**Fig. 3a**). Our group recently demonstrated that in patients hospitalized with SBP, *E. coli* was the predominant isolate in both ascites fluid and bloodstream cultures, with its prevalence increasing with the severity of liver disease ^9^. In a cohort of patients with incident cirrhosis from the VOCAL cohort, we evaluated the hazard of hospitalization with culture-positive peritoneal fluid and/or blood infections in new lactulose initiators (n=12,289) versus non-initiators (n=93,130) using a 1:1 propensity score matching (PSM) approach, with key matching variables including demographics, comorbidities, severity of liver disease, and pharmacotherapies plausibly associated with infection risk, among others (**Fig. 3b**). The primary Cox regression analysis, additionally adjusted for rifaximin use, revealed a significant positive association between lactulose initiation and the overall risk of disseminated infection (hazard ratio [HR] 1.15, 95% CI 1.05-1.26, p=0.002; **Fig. 3c**). Across 500 bootstrap resampled PSM cohorts, only *E. coli* infection hospitalizations were significantly increased in lactulose initiators relative to non-initiators (pooled HR 1.21, 95% CI 1.09-1.33), with *Klebsiella* showing a similar trend, while other species showed no statistically significant differences in risk (i.e., each 95% CI contained 1.00; **Fig. 3d**). The distribution of these estimates in resampled PSM cohorts demonstrated a clear association with *E. coli* and similar trend observed for *K. pneumoniae*, although again this did not reach statistical significance (**Fig. 3e**). This study identifies a previously unrecognized potential side effect of a widely prescribed medication and underscores the need for further consideration of the unintended effects of lactulose therapy on the microbiome.

**Figure 3.**
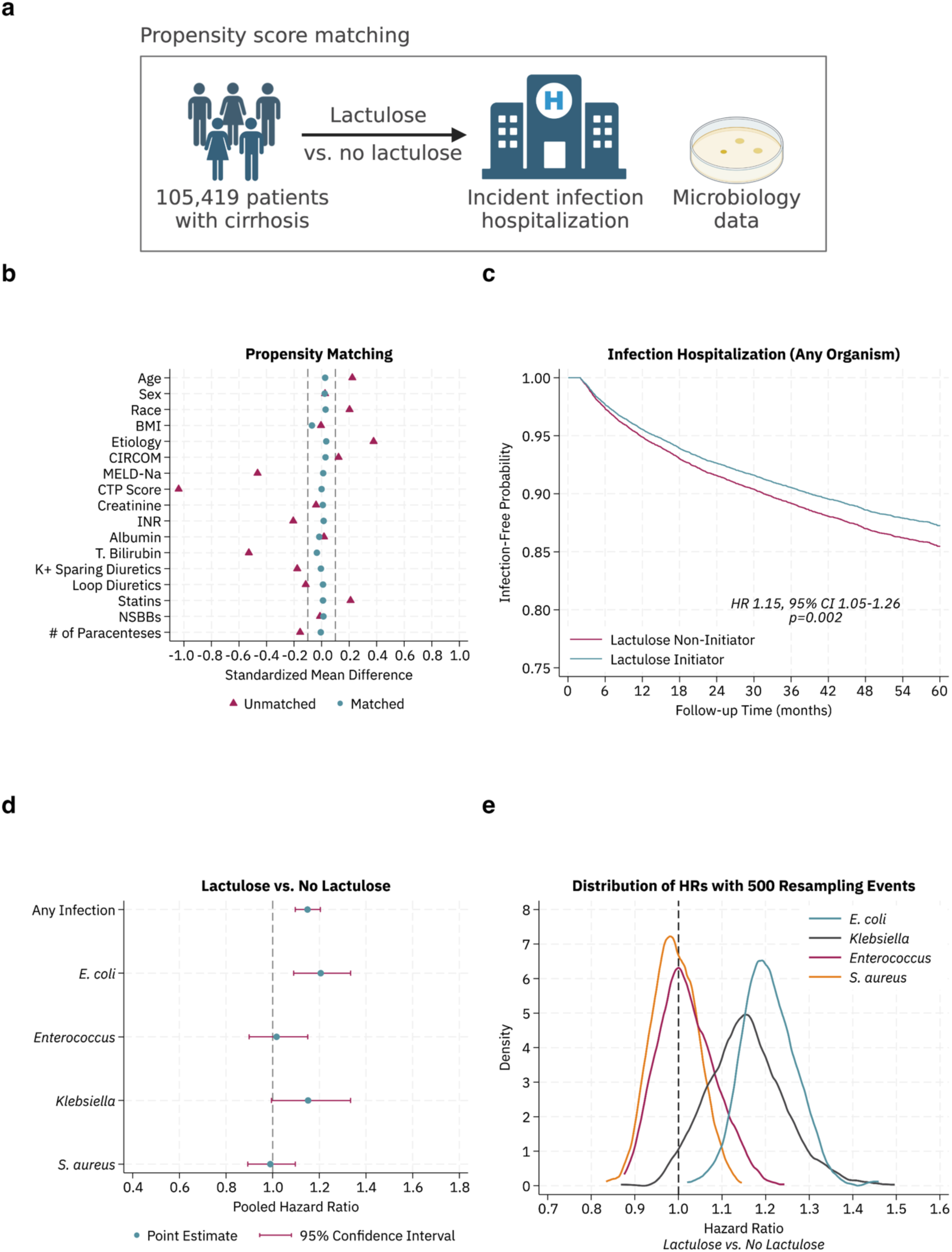
Lactulose increases the risk of *E. coli* disseminated infection in patients with cirrhosis. **A.** Diagram of propensity score matching study design. **B-E**. Results of a retrospective cohort study of patients with incident cirrhosis comparing new initiators of lactulose therapy to controls (N=105,419 patients, 12,289 with lactulose therapy, 93,130 without lactulose therapy). A 1:1 propensity score matching (PSM) algorithm isolated similar patients with and without lactulose exposure, with a standardized mean difference between -0.1 and 0.1 demonstrating excellent covariate balance in the matched cohort (**B**). Cox regression analysis in the matched cohort of infection-free probability (**C**) and pooled hazard ratio of any infection and by bacterial genus/species (**D**) were tested. Distribution of hazard ratios in 500 resampled cohorts (**E**).

### Dietary raffinose suppresses *E. coli* adaptation to lactulose

Given the critical role of lactulose as a therapeutic, we sought to identify strategies to mitigate the unintended side effect of *E. coli* mutation leading to enhanced growth. LacY, the importer of lactulose, is a promiscuous transporter with broad carbohydrate affinity ^37^. We hypothesized that a LacY substrate that is not metabolized by *E. coli* could competitively inhibit both evolution and growth in response to lactulose. Raffinose, a common dietary trisaccharide imported by LacY but generally not utilized by *E. coli*, was selected for testing ^37,38^. Consistent with this, we found that WT *E. coli* had no detectable growth on raffinose over the course of six days, while the LacI^-^ strain exhibited minimal growth (**Fig. 4a**) ^38^. *In vitro* experiments demonstrated that raffinose inhibited the outgrowth of *E. coli* mutants compared to lactulose alone (**Fig. 4b**). To determine whether raffinose could similarly limit the formation of lactulose-metabolizing mutants *in vivo*, mice were colonized with WT *E. coli* and treated with lactulose in the presence or absence of raffinose (**Fig. 4c**). Raffinose co-administration significantly reduced the proportion of *E. coli* capable of metabolizing lactulose (**Fig. 4d**) and blunted the lactulose-induced increase in colonization (**Fig. 4e**).

**Figure 4.**
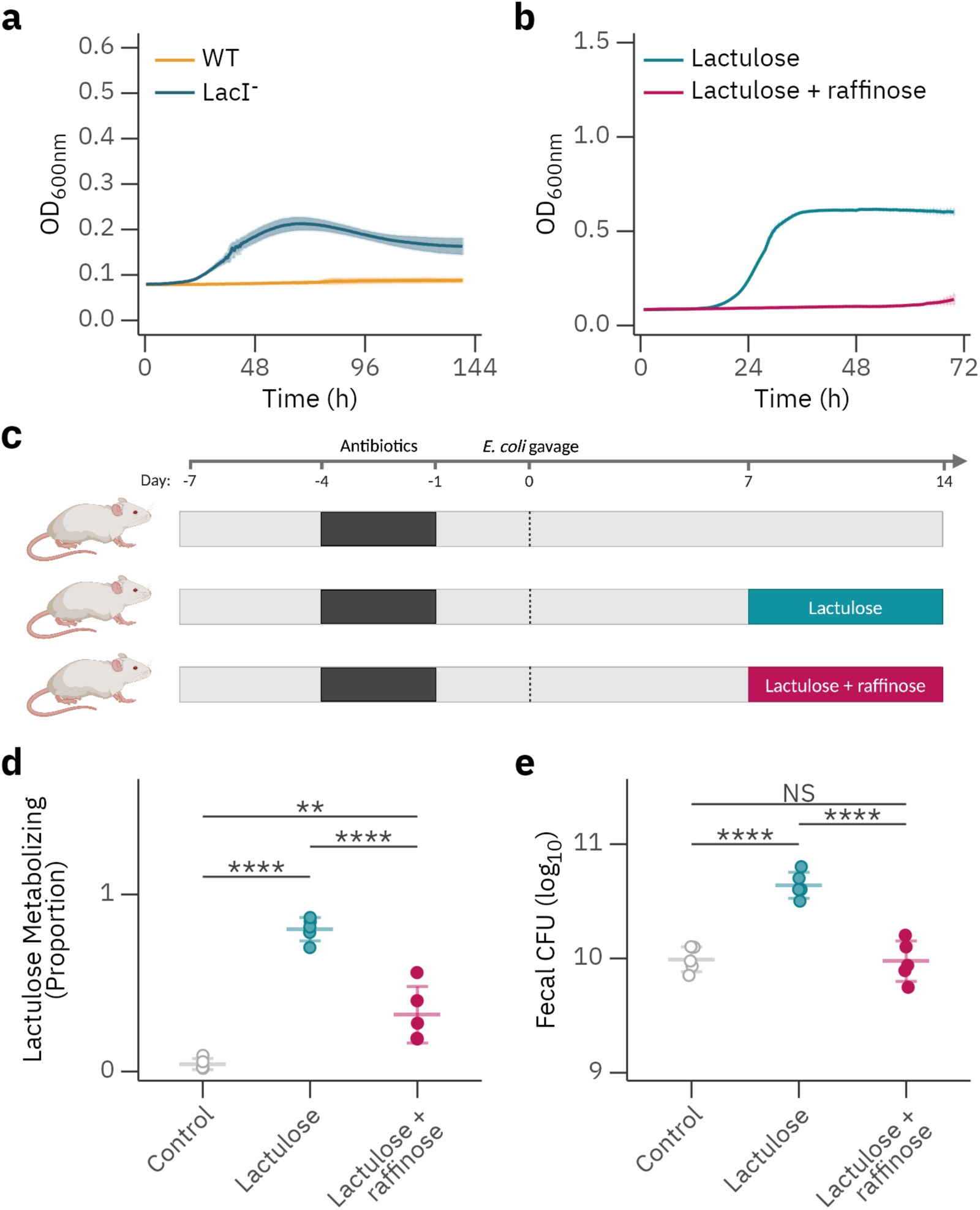
Dietary raffinose inhibits *E. coli* evolution to lactulose while sequential exposure enables a novel metabolic phenotype. **A**. Growth curves of WT and LacI^-^ *E. coli* strains in minimal media containing raffinose (n=3 replicates per condition, results representative of three independent experiments). **B.** Growth curves of WT *E. coli* with lactulose alone or lactulose with raffinose (n=3 wells per group; representative of three independent experiments). **C.** Diagram of murine experimental design. **D and E.** Mice were colonized with WT *E. coli* and treated with lactulose or lactulose and raffinose in the drinking water. Percentage of lactulose-metabolizing mutants (**D**) and fecal CFU (**E**) in indicated experimental groups at day 14 of the study (n=5 mice per group; data representative of two independent experiments). Data shown as mean ± s.d. Statistical analysis was performed using ordinary one-way ANOVA with Bonferroni correction for multiple comparisons (**D and E**). **p<0.01, ****p<0.0001, n.s., not significant.

Together, these findings demonstrate that dietary raffinose can mitigate lactulose-driven colonization of *E. coli* by competitively inhibiting LacY-mediated transport. This highlights the interplay between diet, microbial adaptation, and metabolic evolution, suggesting that strategic carbohydrate interventions could modulate microbiota behavior and reduce pathobiont expansion in the gut.

## Discussion

Here we evaluated the impact of lactulose, a frequently prescribed carbohydrate therapy for cirrhotic liver disease, on gut pathobionts within the microbiota. We demonstrated that lactulose consumption rapidly selects for mutations in the regulatory components of the *E. coli lac* operon across *in vitro*, murine, and clinical contexts. These mutations enhance the metabolic capabilities of *E. coli*, increasing its colonization fitness in the gut. By focusing on lactulose, we identified clear correlations between patient carbohydrate consumption and bacterial phenotypes and genotypes. The alteration of lactulose metabolism by mutations in *lac* operon regulation, described *in vitro*, in mice, and humans, suggests potential relevance to human pathophysiology. This work reveals a potential unanticipated risk of lactulose therapy: the metabolic adaptation of gut pathobionts exploiting an abundant host-consumed carbon source, leading to increased colonization that could be associated with an increased risk for systemic infection.

Lactulose has been a standard of care treatment for HE for over 50 years ^23,24^. Its therapeutic effects are thought to depend on gut microbial metabolism, which reduces ammonia absorption and generates short-chain fatty acids (SCFAs) that suppress gut pathobiont growth. However, our findings offer evidence that suggest a previously unanticipated risk in patients with cirrhosis, where commensal microbes are often depleted, that lactulose may paradoxically enhance the fitness of pathobionts like *E. coli* ^2,3,28^. These results underscore that baseline microbiota composition may play an important role in determining the risks and benefits of lactulose treatment, a factor not currently considered in clinical practice. Given that the mainstay therapies for HE, including lactulose and rifaximin, act through gut microbiota modulation, there is a critical need to consider their effects in contexts where the microbiota is severely disrupted ^24,39^. Future clinical studies should consider incorporating microbiota sequencing as a factor in guiding therapy selection.

Our clinical study has inherent limitations due to its retrospective design, leaving room for residual confounding after propensity score matching. Nevertheless, the specificity of our findings, where *E. coli* dissemination showed a significant signal and *K. pneumoniae* exhibited a weaker trend while *Enterococcus* remained unaffected, reduces the likelihood of systematic confounding. Furthermore, complementary data from murine models and human isolates provide strong biological plausibility for the observed associations. Randomized controlled trials will be required to establish the causal relationship between lactulose therapy and infection risk and to explore the impact of microbiota composition on therapeutic outcomes.

We have demonstrated that constitutive expression of the *lac* operon imposes significant fitness costs during intestinal colonization. This is consistent with previous studies implicating proton motive force disruption via LacY permease transport of non-metabolized substrate ^31^. Interestingly, our murine models demonstrated that loss-of-function mutations in the *lac* operon arise even in the absence of lactulose, suggesting either substrate-independent fitness costs or the presence of unidentified LacY substrates in the gut. These findings were mirrored in human subjects undergoing lactulose therapy, who exhibited spontaneous emergence of such mutations, possibly driven by the intermittent dosing of lactulose. Understanding the evolutionary dynamics of these adaptations, particularly in response to variable exposure, will be an important avenue for future research.

Diet is a well-established driver of microbiota composition and function ^14,17,19,22^, and long-term exposure to dietary compounds can select for novel metabolic capabilities in commensal organisms ^40–42^. Our results reveal an underappreciated force of carbohydrates on shaping the microbiome through rapid evolution. The strong selective pressure exerted by lactulose underscores the carbon-limited environment of the gut for *Enterobacteriaceae* ^22^. Notably, we found that the combination of carbohydrate exposures dictates evolutionary trajectories. This suggests that dietary history could imprint a unique evolutionary signature on the microbiota, potentially contributing to interindividual variability in microbial responses to diet.

Our findings likely extend beyond lactulose and *E. coli*. Dietary selection for metabolic adaptations in commensal bacteria has been previously reported, though the specific substrates driving these changes remain largely unknown ^41,43^. Given the complexity of whole-food carbohydrates and microbiota-derived byproducts, selection for mutants may play a significant role in shaping interpersonal differences in microbiota evolution. Furthermore, our clinical data linking a consumed carbohydrate to infectious dissemination highlights the broader implications of dietary interventions on disease risk. These insights emphasize the need for precision approaches in dietary and therapeutic strategies targeting the microbiota.

## Methods

### Bacterial strains, culture conditions, and antibiotics

Bacterial strains used in these studies are noted in **Supplemental Table 3**. Routine propagation was performed in LB (miller) broth or agar aerobically at 37°C unless otherwise noted. Tetracycline, kanamycin, chloramphenicol, hygromycin, and ampicillin were used at concentrations of 15, 50, 12.5, 100, and 100 μg/mL, respectively.

### Strain construction

The in-frame *lacZ* deletion was constructed by recombineering, as previously described ^44^, except electrocompetent cells of MP1 were prepared by washing with an ice cold solution containing 20% glycerol and 1 mM unbuffered 3-(N-morpholino)propanesulfonic acid (MOPS) ^45^. Electrocompetent cells were electroporated with a PCR fragment obtained from template pKD13 ^44^ and primers lacZ-del-MP1-U and lacZ-del-MP1-L (**Supplemental Tables 4 and 5**). Recombinants were selected on kanamycin and verified by PCR. The Δ*lacZ*::FRT-Kan::FRT was moved into strain MP13, by transduction with P1vir. The kanamycin resistance cassette was removed via Flp recombinase by transformation with pCP20 as described ^44^.

For complementation of the LacI^-^ phenotype, the MP13 LacI^-^ *E. coli* strain was transformed with pTrc99a, which expresses the LacI protein, via electroporation ^46^.

To mark the human isolate 36-1, chromosomal insertion of kanamycin resistance was accomplished using the pUC18R6KT-mini-Tn7T-Km plasmid ^47^, which was delivered via conjugation from an *E. coli* MFDpir. To mark the human isolate 36-2 with chloramphenicol resistance, the kanamycin resistance cassette was replaced from plasmid pUC18R6KT-mini-Tn7T-Km with a chloramphenicol resistance cassette. This was generated via FLP recombinase, expressed on pFLP-hyg and a chloramphenicol resistance cassette flanked by FRT-sites, amplified from pKD3 with primers CAM-FRT-NotI-u1 and CAM-FRT-ClaI-l1 ^48,49^. This PCR product was electroporated into competent PIR2 *E. coli*. This generated plasmid pUC18R6KT-mini-Tn7T-Cat. The final 36-2 strain with chloramphenicol resistance was constructed via diparental mating with an MFDpir strain containing pUC18R6KT-mini-Tn7T-Cam^50^. Insertion into the correct chromosomal site was confirmed via colony PCR using previously described primers ^48^.

### Growth curves

To determine the growth characteristics of bacterial strains, overnight cultures of bacterial isolates were pelleted and washed in PBS and diluted 1:1,000 in M9 minimal media supplemented with trace elements containing carbon sources as indicated at a final concentration of 20mM. OD_600_ was determined via the a BioTek Epoch2 plate reader and grown with continuous shaking at 37°C. TDG was added at a concentration of 5mM, where indicated. For human isolate characterization, growth time to mid-log phase on lactulose was determined by the first time point with OD_600_ >0.5.

For serial passage of pediatric bacteremia isolates, strains were grown in M9 minimal medial containing lactulose for 48 hours, after which they were spun down, washed and diluted 1:100 into LB and grown for 24 hours. Finally, the strains were re-tested for growth in M9 lactulose.

### Human subject study and fecal bacterial isolates

The human stool samples used in this study are from baseline samples of patients enrolled in the Acute-on-Chronic Liver Failure with Gut Microbiota-Targeted Assessment and Treatment (ACCLIMATE) study, as previously published ^28^. Stool samples with 5% or greater relative abundance *E. coli* and *K. pneumoniae*, based on previously published shotgun sequencing data ^28^, were selected for bacterial isolation. Fecal samples were resuspended in PBS at a ratio of 1μL PBS: 5mg stool and serially diluted 1:10 in PBS. Serial dilutions were spread on MacConkey agar containing lactose. Colonies were scored for lactose and lactulose utilization using MacConkey agar containing lactose or lactulose at a concentration of 10g/L. 10-12 single colonies (pink or white) were selected at random from the MacConkey lactose plates, restreaked for purity, and stored for further analysis.

### RNA isolation and qPCR

1 ml of mid-exponential culture (OD 0.25-0.3) was pelleted by centrifugation at 8,000xg for 5 minutes and washed twice with PBS. The pellet was resuspended in 700μL of Trizol reagent (Invitrogen) and processed using the Direct-zol RNA Miniprep kit (Zymo Research) following the kit protocol. The resulting RNA was treated with the TURBO DNA-free Kit (Invitrogen) per kit protocol. The purified RNA was converted to cDNA using the High-Capacity RNA-to cDNA Kit (Invitrogen). The cDNA was diluted 1:10 with molecular grade water and used for SYBR Green based qPCR with primers specific to the desired targets and normalized to expression of the house keeping gene gyrA.

### DNA extraction, whole genome sequencing, and genome assembly

Bacterial isolate DNA extraction and WGS were performed by SeqCenter in Pittsburgh, PA. DNA extraction was performed with ZymoBIOMICS ™ DNA Miniprep Kit2. 50-100mg of plated isolates were scraped and resuspended in 750 µl of lysis solution. Samples were transferred to ZR BashingBead™ Lysis Tubes and mechanically lysed using the MP FastPrep-24™ lysis system with 1 minute of lysis at maximum speed and 3 minutes of rest for 2 cycles. Samples were then centrifuged at 10,000rcf for 1 minute. 400µl of supernatant was transferred from the ZR BashingBead™ Lysis Tube to a Zymo-Spin™ III-F Filter and centrifuged at 8,000rcf for 1 minute. 1,200 µl of ZymoBIOMICS™ DNA Binding Buffer was added to the effluent and mixed via pipetting. 800µl of this solution was transferred to a Zymo-Spin™ IICR Column and centrifuged at 10,000rcf for 1 minute. This step was repeated until all material was loaded onto the Zymo-Spin™ IICR Column. DNA bound to the Zymo-Spin™ IICR Column was washed 3 times with 400µl and 700µl of ZymoBIOMICS™ DNA Wash Buffer 1 and then 200 µl of ZymoBIOMICS™ DNA Wash Buffer 2 with a 1-minute spin down at 10,000rcf for each, respectively. Washed DNA was eluted using 75µl of ZymoBIOMICS™ DNase/RNase Free Water following a 5-minute incubation at room temperature and a 1-minute spin down at 10,000rcf. The Zymo-Spin™ III-HRC Filter was prepared using 600 µl of the ZymoBIOMICS™ HRC Prep Solution and a centrifugation at 8,000rcf for 3 minutes. Eluted DNA was then purified by running the effluent through the prepared Zymo-Spin™ III-HRC Filter. Final DNA concentrations were determined via Qubit.

Illumina sequencing libraries were prepared using the fragmentation-based and PCR-based Illumina DNA Prep kit and custom IDT 10bp unique dual indices (UDI) with a target insert size of 320 bp. No additional DNA fragmentation or size selection steps were performed. Illumina sequencing was performed on an Illumina NovaSeq 6000 sequencer in one or more multiplexed shared-flow-cell runs, producing 2×151bp paired-end reads. Demultiplexing, quality control and adapter trimming was performed with bcl-convert (v4.1.5). De novo genome assembly was performed using Shovill (v1.1.0) (https://github.com/tseemann/shovill).

### Phylogenetic tree and SNP analysis

The assembled genomes were annotated using bakta (v1.7.0), using default parameters ^51^. Panaroo (v1.2.10) with clean mode set as strict and with mafft (v7.508) as the aligner, was then used to construct the core gene pangenome for the 18 genomes identified as *E.coli* in the ACCLIMATE dataset ^52^. The core threshold was set to 0.98. Subsequently, a maximum-likelihood phylogenetic tree with 1000 bootstrap replicates and the model set to HKY was constructed using iq-tree (v 2.2.0.3) ^53^. Using the tool mlst (v2.22.1),13 different sequence types were identified in the 18 study isolates ^54^. The tree was visualized and annotated using iTol ^55^.

To determine the number of SNPs between genomes 36-1 and 36-2, snippy (v4.6.0) was used with 36-1 as the reference ^56^. This analysis revealed 57 SNPs between the two genomes.

### Beta-galactosidase assay

Overnight cultures of relevant strains were diluted 1:100 in LB with or without IPTG (1mM) or lactulose (20mM). Cultures were grown to early exponential phase at OD_600_ ∼0.3. Cultures were pelleted in a 96-well plate and resuspended in 200µL B-PER™ (Thermo Scientific) supplemented with lysozyme (10µg/mL). Resuspended cells were allowed to lyse in this mixture for 20 minutes at room temperature. BG assay mix was made from 1x BG assay salts with 0.1mg/mL ortho-Nitrophenyl-β-galactoside (ONPG) (10x BG assay salts: 600 mM Na_2_HPO_4_, 400 mM NaH_2_PO_4_, 100 mM KCl, 10 mM Mg SO_4_). 120µL BG assay mix was added to 80µL lysed culture in a fresh 96-well plate. OD_420_ was read every 1 minute for 30 minutes with a BioTek Epoch2 plate reader. Relative quantification of BG activity was determined by dividing the linear slope of the OD_420_ curve by the terminal OD_600_.

### Genetic locus comparison

For sequencing of the *lac* operon and *lacI* from mouse fecal isolates, the primers lac-cPCR-u1 and lac-check-l1 were used to amplify this region from MP1 and 36-2 via PCR (**Supplemental Table 5**). Linear/PCR sequencing was performed by Plasmidsaurus using Oxford Nanopore Technology with custom analysis and annotation. Alignment and visualization were performed using Geneious Prime 2024.0.7.

### Mice

SPF C57Bl/6J female mice were purchased from The Jackson Laboratory at 8 weeks of age. Mice were housed under standard lighting cycle conditions (12 hours on/12 hours off) and provided acidified water. There was no investigator blinding for these studies, and no animals were excluded from analysis.

### Mouse colonization model

For *E. coli* colonization, mice were placed on a fiber-free diet (TestDiet 5Z6G) 1 week prior to initiation of the study, as previously published ^22^. Mice were pre-treated with vancomycin (0.5g/L) and aspartame (5g/L) in the drinking water for 3 days. 24 hours after antibiotic treatment, mice were orogastrically gavaged with 10^8^ CFU of *E. coli* in 100μL, as indicated. Inoculum was generated from aerobically grown overnight cultures in LB Miller media and then washed and diluted 1:10 in sterile PBS. Fecal samples for CFU analysis were collected, homogenized in PBS, serially diluted, and spot-plated on LB agar supplemented with tetracycline. Plates were incubated for 12 hours at 37°C, colonies were counted, and CFU were calculated per weight of fecal sample. Prior to and after antibiotic treatment, but before gavage, each mouse was routinely tested for resistant colonies. For lactulose treatment, mice were provided lactulose in the drinking water at 5% (weight/volume). For combined treatment with raffinose, this was provided in the drinking water at a 5:1 molar ratio with lactulose.

### *In vitro* and *in vivo* strain competitions

To differentiate between strain derivatives of MP1 during mixed competitions in culture or murine models, we used a previously published system of fluorescent marker-based quantification ^57^. In brief, strains were differentially marked with chromosomally encoded tetracycline-inducible GFP or mCherry. Serial dilutions of culture or stool inoculated with competing strains were plated on tetracycline-containing LB and grown for 24 hours. Fluorescent colonies were quantified, and competitive index was calculated as the output ratio divided by the input ratio of the competing strains.

For differentiation of strains 36-1 and 36-2, chromosomally encoded kanamycin and chloramphenicol cassettes enabled differential growth on LB agar containing the appropriate antibiotic. Quantification of each strain was performed from culture or mouse stool via homogenized, serially dilution, and plating on two separate plates containing these antibiotics. For lactulose metabolism phenotyping, mouse stool was plated on MacConkey agar with lactulose containing chloramphenicol to select for strain 36-2.

### Mouse fecal isolate characterization

For strain characterization from mouse stool, five individual colonies were isolated per mouse from fecal samples plated on LB agar with tetracycline. These were restreaked for purity, and stored for further analysis. Each strain was subjected to spot plating on MacConkey lactose or lactulose, growth evaluation in lactulose, and BG activity testing. Selected amplicon sequencing was performed on selected strains as indicated.

### Epidemiology studies

#### Study design and cohort selection

To explore the association between new lactulose initiation and infection hospitalization in patients with cirrhosis in a clinical cohort, we performed a retrospective cohort study using data from a well-established Veterans Health Administration (VHA) dataset called the Veterans Costs and Outcomes Associated with Liver Disease (VOCAL) ^58^. The VOCAL dataset contains granular longitudinal data from over 100,000 patients with cirrhosis across 128 VHA centers from 1/1/2008 through 12/31/2023 and has been utilized for numerous natural history studies of patients with liver disease ^36,59,60^. For this study, we identified patients >18 years old with incident cirrhosis diagnosis, which was determined using a validated VHA algorithm based on International Classification of Diseases (ICD)-9/10 codes ^61,62^. Patients were excluded if they developed an infection-related hospitalization within two months of lactulose initiation, after cirrhosis diagnosis (i.e., two-month lag). This exclusion was applied to mitigate the possibility of confounding by indication, e.g., an infection was already present or developing that drove initiation of lactulose. Patients were also excluded if they were ever exposed to lactulose prior to the new initiation window around the time of cirrhosis diagnosis.

#### Exposures and outcomes

For each patient we collected detailed data on demographics (age, sex, race/ethnicity), body mass index, cirrhosis comorbidity index (CIRCOM score), medication data including potassium-sparing diuretics, loop diuretics, non-selective beta blockers, and statins, number of paracenteses in the 60 days prior to cirrhosis diagnosis, and lab data at the time of cirrhosis diagnosis (creatinine, albumin, bilirubin, INR, sodium); these lab data were used to calculate model for end-stage liver disease-sodium (MELD-Na). Etiology of liver disease and Child-Turcotte-Pugh (CTP) score and class were classified using validated VHA algorithms ^58,63,64^. The primary exposure of interest was new initiation of lactulose, defined as first exposure within 60 days of cirrhosis diagnosis. This was ascertained using the VHA inpatient and outpatient pharmacy tables, which provides comprehensive information on prescription fills across all VHA sites. Rifaximin new initiation was ascertained in similar fashion.

The primary outcome was time to hospitalization with positive bacterial culture on inpatient blood or peritoneal fluid sampling. Secondary outcomes included separate hospitalization infection with positive bacterial cultures (blood or peritoneal) from the following microorganisms of interest: *E. coli*, *Klebsiella*, *Enterococcus*, *S. aureus*. These data were obtained from VHA microbiology results tables.

## Statistics

GraphPad Prism (version 10.2.1) was used to perform statistical analysis for culture and mouse studies. For comparisons of two groups, two-tailed Student’s t-test was used. For comparisons of three groups or more, ordinary one-way ANOVA was performed with Bonferroni correction for multiple comparisons. For longitudinal comparison of two groups, multiple t-tests were performed with Holm-Sidak correction for multiple comparisons.

For epidemiology studies, to evaluate the association between lactulose and risk of infection hospitalization a 1:1 propensity matching (PSM) approach was used to achieve covariate balance in new lactulose initiators versus non-initiators. This entailed fitting a multivariable logistic regression model containing the following key *a priori* baseline covariates: age, sex, race/ethnicity, body mass index, etiology of liver disease, CIRCOM, MELD-Na, CTP score, creatinine, albumin, bilirubin, INR, potassium-sparing diuretics, loop diuretics, statins, non-selective beta blockers, and number of paracenteses. A propensity score was generated from this model and a 1:1 nearest neighbor matching approach was used with a caliper width of 0.01. To assess for post-matching covariate balance we then computed standardized mean differences (SMDs) for each variable. An SMD within +/-0.1 was regarded to reflect excellent balance, consistent with best practice recommendations ^65^. In the propensity matched cohort, Cox regression analysis was used to evaluate the relative hazard of infection hospitalization with lactulose exposure. Time zero was cirrhosis diagnosis and patients were right-censored at maximum follow over a 60-month time horizon. The Cox model was adjusted for rifaximin exposure (note this was not included in the propensity score due to lactulose collinearity). A hazard ratio and 95% confidence interval for any infection hospitalization were presented for lactulose exposure and adjusted survival curves were plotted. To ensure robustness of findings, bootstrapping was performed where 500 random seed iterations of propensity score generation were used to estimate pooled relative hazards of any infection, as well as specific microorganism infections. The distribution of hazard ratios were displayed using kernel density plots for any infection, *E. coli* infection, *Klebsiella* infection, *Enterococcus* infection, and *S. aureus* infection, and in pooled fashion using coefficient plots. Structured query languages (SQL) and STATA/SE 18.0 were used for data management and analyses in epidemiology studies.

## Study approval

All research has been conducted in an ethical and responsible manner, and is in full compliance with all relevant codes of experimentation and legislation. All animal studies were conducted in accordance with ethics regulations under protocols approved by the University of Pennsylvania IACUC and Biosafety Committees. The ACCLIMATE study was performed under formal approval of the Institutional Review Board of the University of Pennsylvania (IRB protocol #827492). All patients were consented prior to inclusion in the study. Pediatric bacteremia isolates were collected under CHOP IRB protocol #14648. Bacteremia isolates were completely deidentified for the study with no patient information associated. The epidemiologic studies received IRB approval from the Corporal Michael J. Crescenz Philadelphia Veterans Affairs Medical Center.

## Data availability

Whole genome sequencing and genetic locus sequencing data are available under BioProject number PRJNA1232022.

## Author contributions

A.L.H., M.G., and G.D.W. conceived and planned the overall study design and supervised the project. A.L.H., S.C., J.C., J.L., and L.H. performed the murine studies. A.L.H., S.C., J.Y.C., G.P.B. and E.S.F. performed the *in vitro* studies. A.L.H. isolated and tested bacterial mutants, and human and murine fecal bacterial clones. A.L.H. and M.R. constructed the bacterial strains. E.T. and A.M.M. created the phylogenetic tree and performed SNP analysis. B.E.G. and S.J.M collected and processed the bacteremia isolates. A.M.M., P.J.P., and J.P.Z. collated and provided access to the bacteremia isolates and provided bacterial genomics insights. D.E.K., M.S., and N.M. provided epidemiologic analysis. G.D.W. and R.R. provided critical clinical insight and access to clinical biospecimens. A.L.H., M.G., and G.D.W. wrote and edited the manuscript with input from all the authors.

## Acknowledgements

This work was supported by the Division of Gastroenterology and Hepatology at the Hospital of the University of Pennsylvania and the CHOP Microbiome Center. A.L.H. is funded by the NIH (K08DK138319), the American Gastroenterological Association Research Scholar Award (AGA2024-13-02) and Crohn’s and Colitis Foundation Research Fellows Award (1049281). N.M. is funded by the NIH (K08-DK124577, R03-DK134794) and through investigator-initiated funding (Grifols) unrelated to this work. M.G. is funded by NIH R35GM139541. This work is also supported by the Sherman Prize (to G.D.W.) and the Penn Dean’s Innovation Fund Award (to G.D.W.). J.Y.C. was Funded by the AGA Research Foundation’s AGA-Dr. Harvey Young Education and Development Foundation’s Young Guts Scholar Program - AGA2025-51-03. P.J.P, A.M.M, and J.P.Z. were supported by the Center for Microbial Medicine at CHOP. We also acknowledge the assistance of the PennCHOP Microbiome Program. We thank the microbial ARchive and Cryocollection (microbialARC) at CHOP. We thank the Microbial Culture and Metabolomics Core of the PennCHOP Microbiome Program, the Center for Molecular Studies in Digestive and Liver Diseases (NIH P30DK050306), the Penn Host Microbial Analytic and Repository Core (H-MARC), and the Penn Center for Nutritional Science and Medicine (PenNSAM) for their technical expertise. We thank Katlyn Bose for assistance constructing *ΔlacZ E. coli* strain MP330. We additionally thank Nathania Hau for contributions in figure visualization and construction.

## Notes

### Competing Interest Statement

The authors have declared no competing interest.

