## Extended Data Figures 1-7 for "Carbohydrate consumption drives adaptive mutations in *Escherichia coli* associated with increased risk for systemic infection"

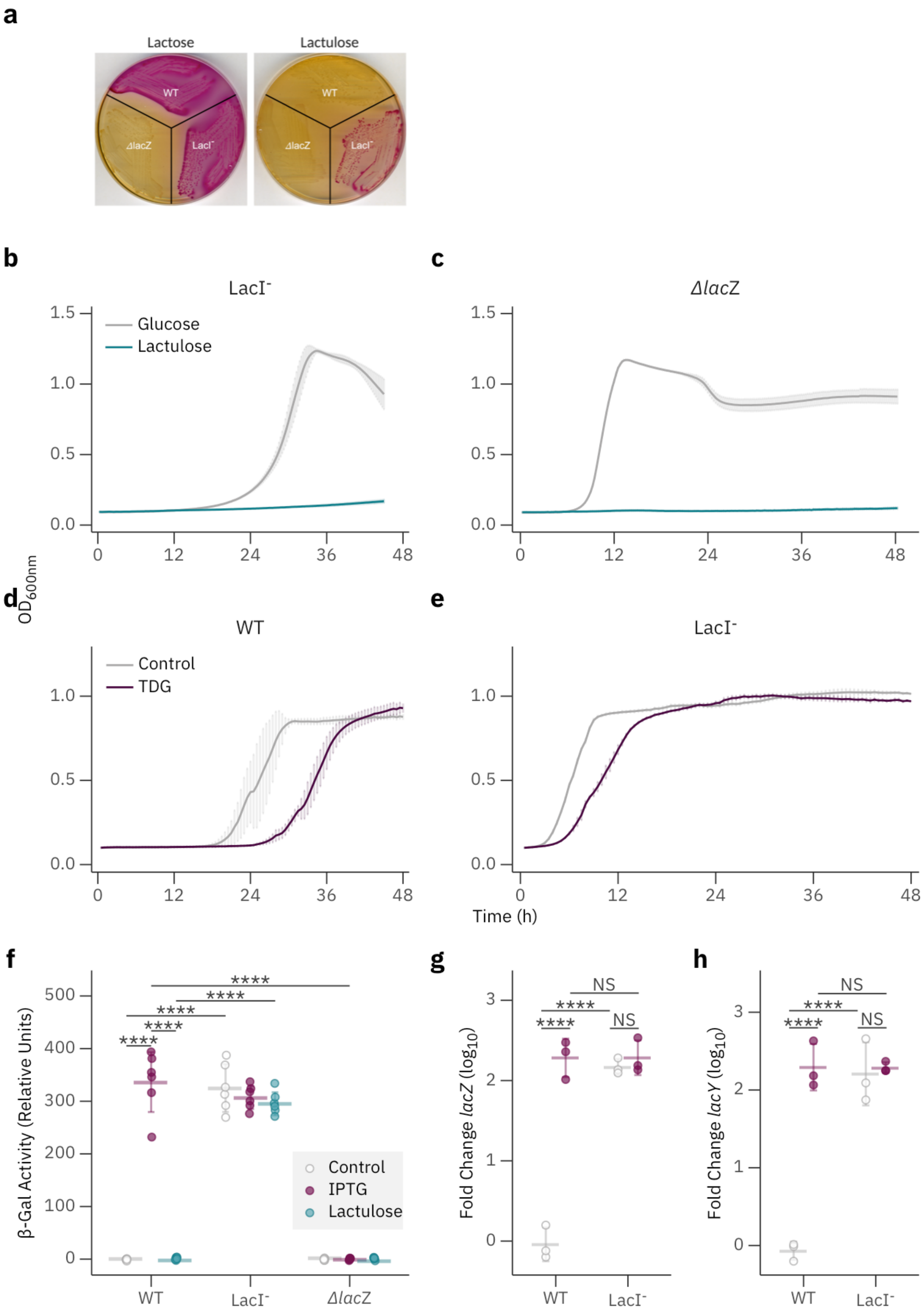
a
Lactose
Lactulose
WT
ΔlacZ
LacI-
b
LacI-
Glucose
Lactulose
c
ΔlacZ
OD600nm
d
WT
Control
TDG
e
LacI-
Time (h)
f
β-Gal Activity (Relative Units)
****
Control
IPTG
Lactulose
WT
LacI-
ΔlacZ
g
Fold Change lacZ (log10)
NS
WT
LacI-
h
Fold Change lacY (log10)
NS
WT
LacI-

**Extended Data Figure 1. A.** Bacterial growth of WT, LacI$^-$ and *ΔlacZ E. coli* strains on MacConkey with lactose or lactulose. Pink color indicates fermentation of the carbohydrate. **B and C.** Growth of LacI$^-$ strain complemented with LacI expressed from a plasmid (**B**) or the *ΔlacZ* strain (**C**) on glucose or lactulose (n=3 wells per condition, results representative of two independent experiments). **D and E.** Growth of WT (**D**) and LacI$^-$ *E. coli* in lactulose with or without β-d-galactopyranosyl 1-thio-β-d-galactopyranoside (TDG) supplemented at a concentration of 5mM (n=3 wells per condition, results representative of three independent experiments). **F.** Beta-galactosidase activity of the three strains noted, grown without IPTG or lactulose (n=6 per group; results are representative of three independent experiments). **G and H.** RNA abundance of *lacZ* (**G**) and *lacY* (**H**) of WT and LacI$^-$ *E. coli* with or without IPTG relative normalized to the WT strain without IPTG condition (n=3 biological replicates per condition). Data shown as mean ± s.d. Statistical analysis was performed using one-way ANOVA with Bonferroni correction (**F-H**), ****p<0.0001.

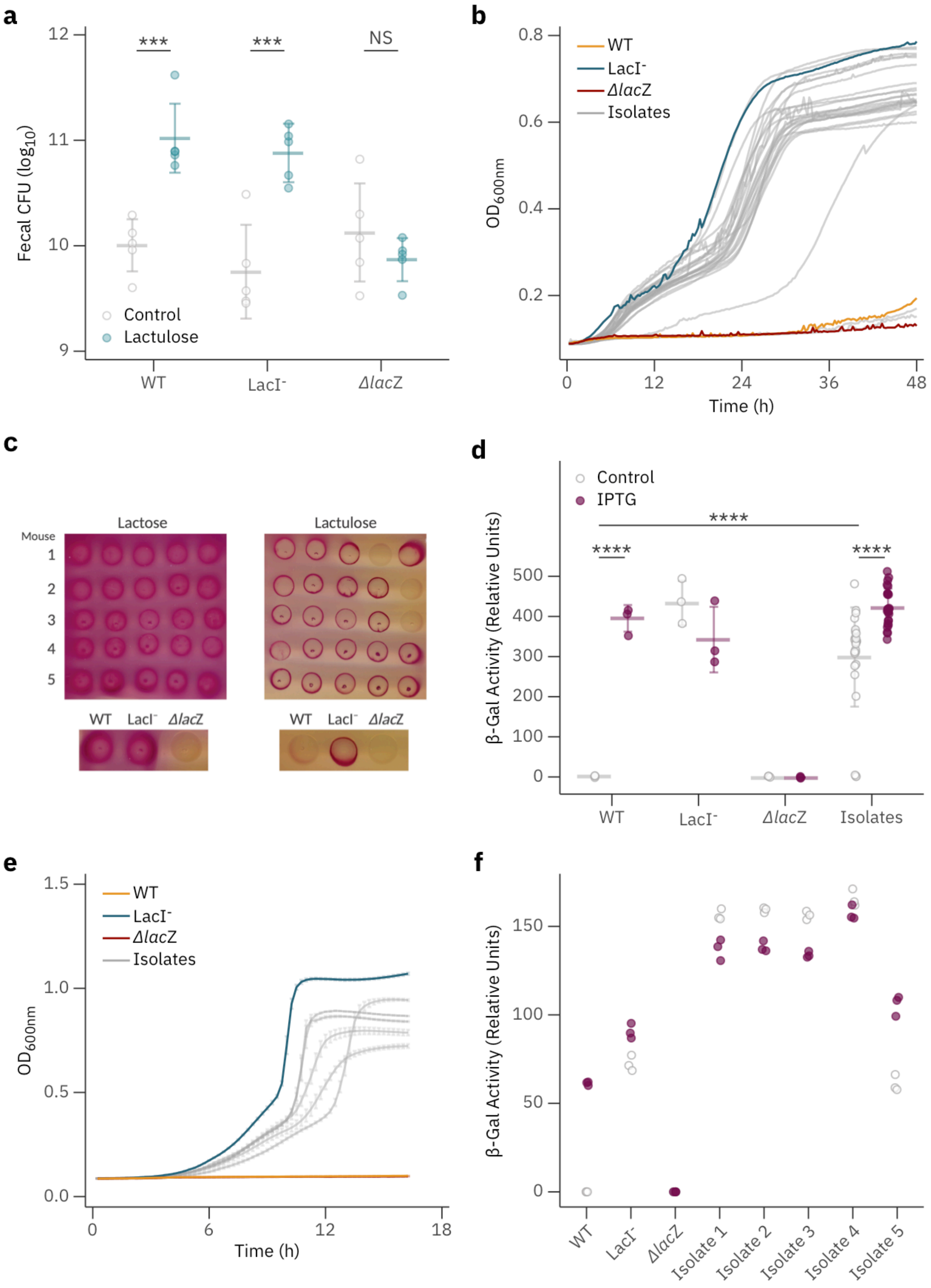
a
Fecal CFU (log10)
12
11
10
9
***
***
NS
Control
Lactulose
WT
LacI-
ΔlacZ
b
0.8
0.6
0.4
0.2
OD600nm
WT
LacI-
ΔlacZ
Isolates
0
12
24
36
48
Time (h)
c
Lactose
Lactulose
Mouse
1
2
3
4
5
WT
LacI-
ΔlacZ
d
Control
IPTG
****
****
****
β-Gal Activity (Relative Units)
500
400
300
200
100
0
WT
LacI-
ΔlacZ
Isolates
e
1.5
1.0
0.5
0.0
OD600nm
WT
LacI-
ΔlacZ
Isolates
0
6
12
18
Time (h)
f
150
100
50
0
β-Gal Activity (Relative Units)
WT
LacI-
ΔlacZ
Isolate 1
Isolate 2
Isolate 3
Isolate 4
Isolate 5

**Extended Data Figure 2. A.** Fecal CFU per gram of mice colonized with WT, LacI$^-$ and *ΔlacZ E. coli* strains after 2 weeks of treatment with lactulose or control (n=5 mice per group, results representative of 2-3 independent experiments). **B-D.** Fecal *E. coli* were isolated from mice colonized with WT *E. coli* and subsequently treated with lactulose in the drinking water for two weeks. Growth curves of individual isolates and WT, LacI$^-$ and *ΔlacZ E. coli* controls in minimal media with lactulose (**B**) (n=1 clone per well, results representative of two independent experiments). Growth of isolates and noted controls on MacConkey agar with lactose or lactulose (**C**) (results representative of two independent experiments). BG activity of isolates and controls with or without IPTG inducer (**D**) (n=25 isolates or n=3 for controls; results representative of two independent experiments). **E and F.** Single isolates (one per mouse) from **B-D** were carried forward for further analysis. Growth curves in lactulose minimal media (**E**) and BG activity without (grey) or with (maroon) IPTG (**F**) were measured. Data shown as mean ± s.d. Statistical analysis was performed using ordinary one-way ANOVA with Bonferroni correction for multiple comparisons (**A and D**). ***p<0.001, ****p<0.0001, n.s. not significant.

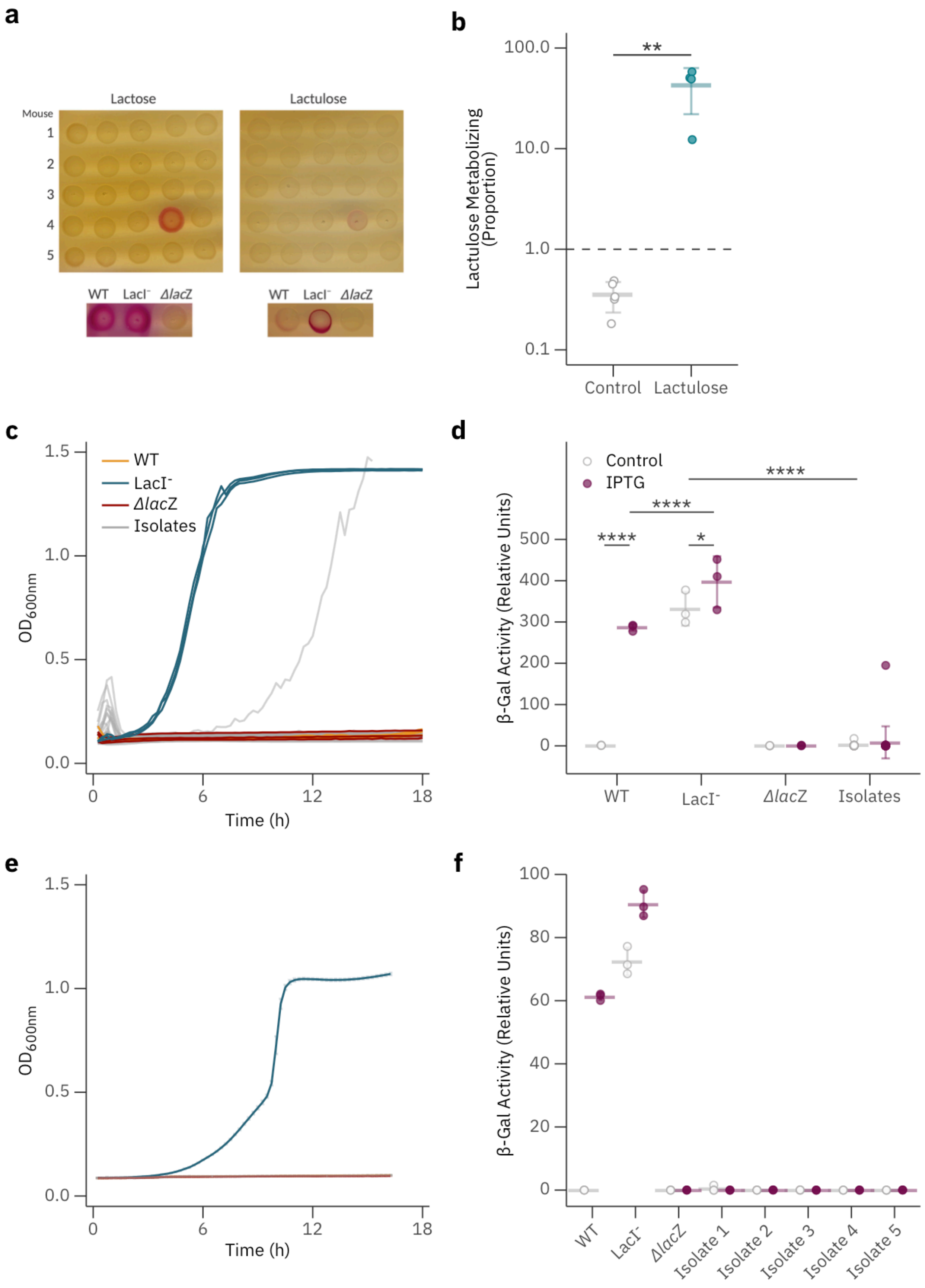
a
Lactose
Lactulose
Mouse
1
2
3
4
5
WT
LacI⁻
ΔlacZ
b
**
100.0
10.0
1.0
0.1
Lactulose Metabolizing (Proportion)
Control
Lactulose
c
WT
LacI⁻
ΔlacZ
Isolates
OD600nm
Time (h)
d
Control
IPTG
****
****
****
*
β-Gal Activity (Relative Units)
WT
LacI⁻
ΔlacZ
Isolates
e
OD600nm
Time (h)
f
β-Gal Activity (Relative Units)
WT
LacI⁻
ΔlacZ
Isolate 1
Isolate 2
Isolate 3
Isolate 4
Isolate 5

**Extended Data Figure 3. A-D.** *E. coli* clones were isolates from feces of mice colonized with $LacI^-$ *E. coli* after two weeks with or without lactulose treatment in the drinking water. Isolates from mice not treated with lactulose were plated on MacConkey with lactose or lactulose (**A**). Control WT, $LacI^-$ and ***Δ****lacZ E. coli* strains are shown for reference (images of controls duplicated from **Supplemental Figure 2C** for comparison; results representative of two independent experiments). Percentage of lactulose metabolizing *E. coli* clones isolated from mice colonized with the $LacI^-$ strain after 2 weeks of treatment with or without lactulose (**B**) (n=5 mice per group, results representative of two independent experiments). Growth curves of isolates and controls in minimal media with lactulose (**C**) (n=1 clone per well, results representative of two independent experiments). BG activity of isolates with or without IPTG inducer and controls (**D**) (n=25 isolates or n=3 per control; results representative of two independent experiments. Majority of data points of isolates are overlapping). **E and F**. Single representative isolates per mouse were selected from **A-D** for further analysis via growth curves in minimal media with lactulose (**E**) and BG activity without (grey) or with (maroon) IPTG (**F**) (n=3 per isolate or control, results representative of two independent experiments). Data shown as mean ± s.d. Statistical analysis was performed using student's t-test (**B**) or ordinary one-way ANOVA with Bonferroni correction for multiple comparisons (**D**). *$p<0.05$, **$p<0.01$, ****$p<0.0001$.

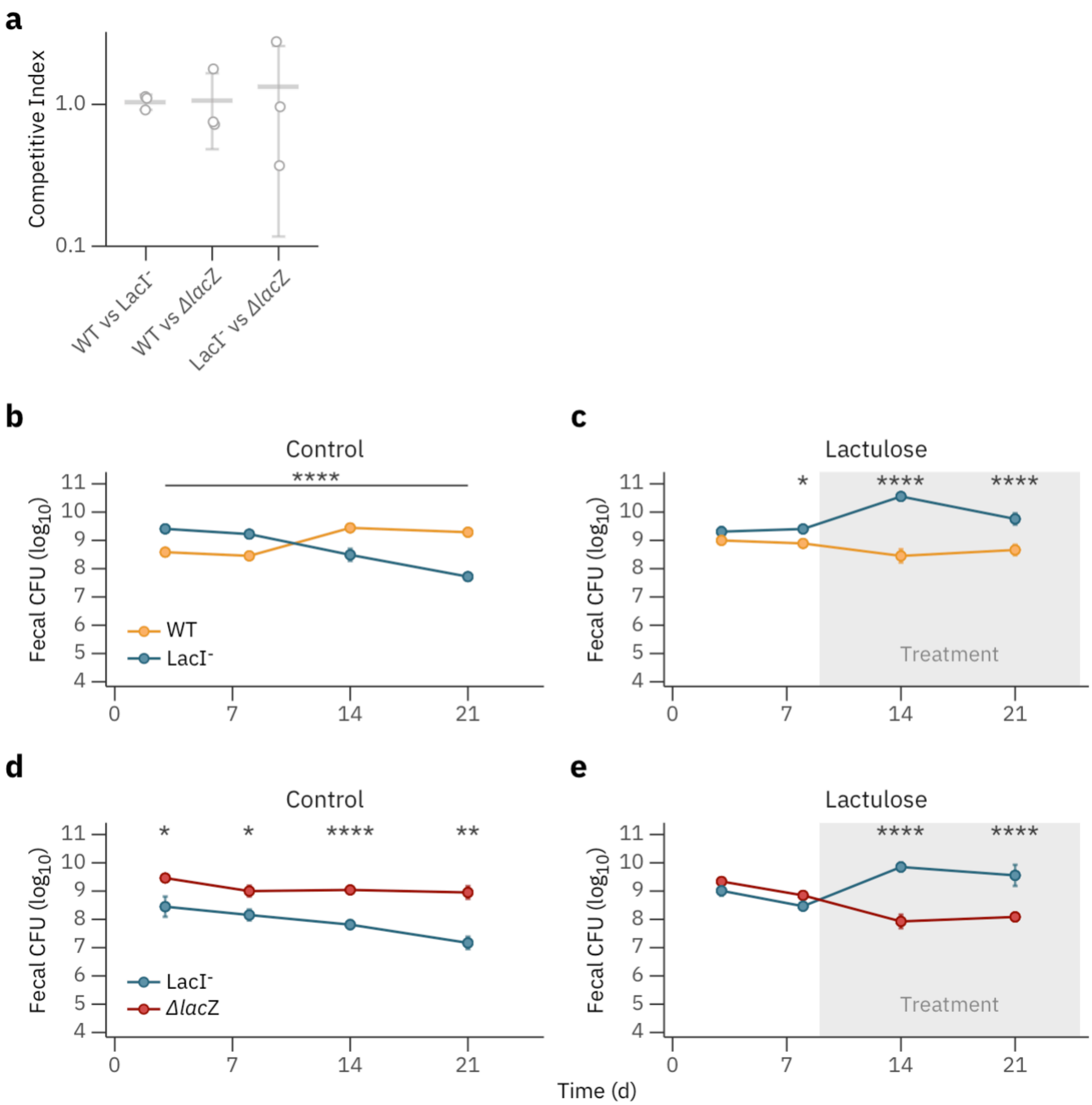


**Extended Data Figure 4. A.** Competitive index of pairwise competitions of WT, LacI- and *ΔlacZ* *E. coli* strains in LB media after 24 hours of growth (n=3 biological replicates per condition, results representative of two independent experiments). **B-E.** Fecal CFU of differentially marked *E. coli* strains in pairwise competitions without (**B and D**) or with (**C and E**) lactulose treatment (n=5 mice per group, results representative of two independent experiments). Time of lactulose initiation is denoted by the grey section. Data shown as mean ± s.d. Statistical analysis was performed using multiple t-tests with Holm-Sidak correction for multiple comparisons (**B-E**). *p<0.05, **p<0.01, ****p<0.0001.

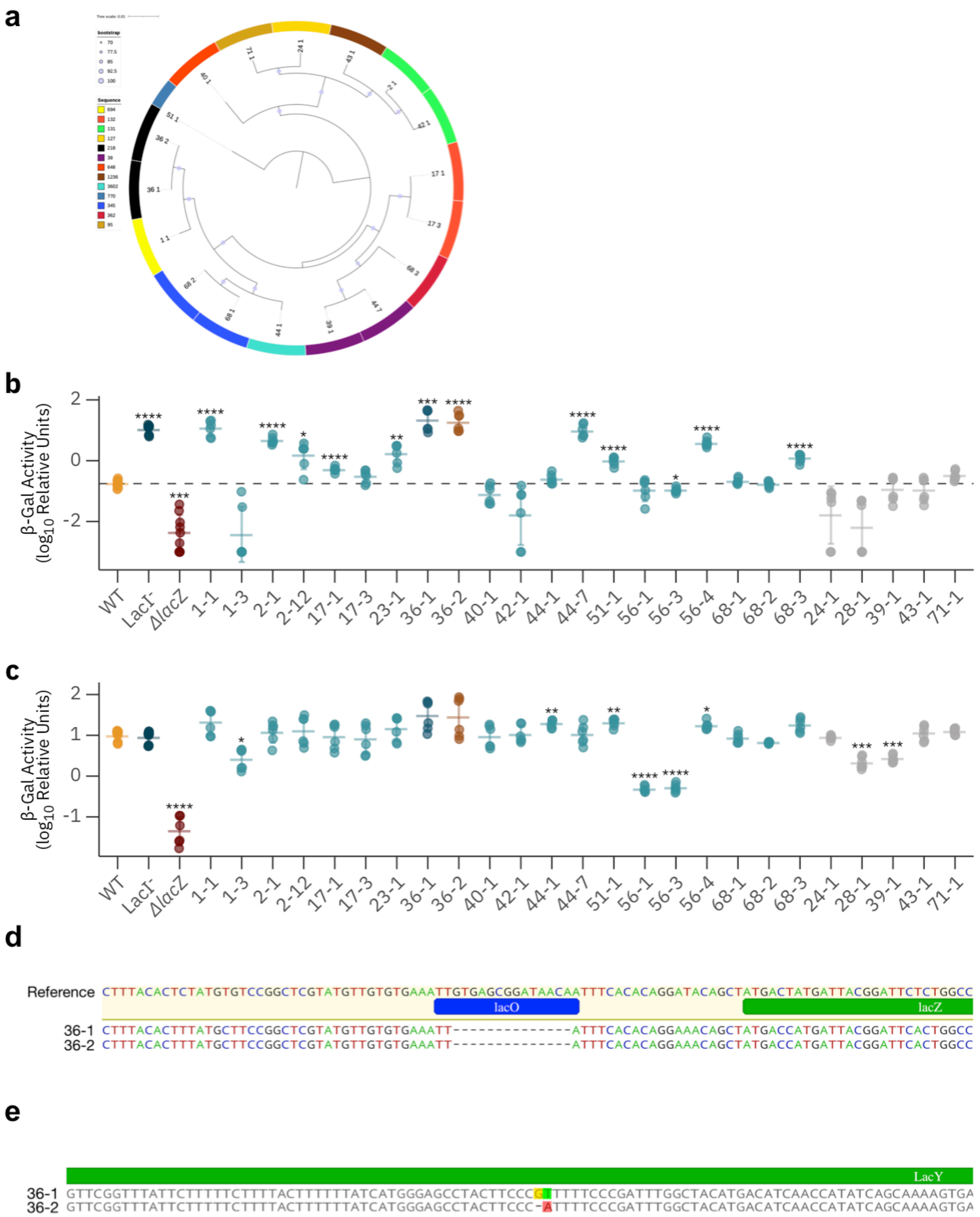


**Extended Data Figure 5. A.** Phylogenetic tree of human fecal *E. coli* isolates. Sequence type noted by color in outside ring. **B and C.** BG activity of human fecal isolates with reference MP13 WT, LacI⁻ and *ΔlacZ E. coli* strain controls without (**B**) or with (**C**) IPTG inducer. Strains isolated from patients on lactulose therapy are noted in light blue, strains 36-1 and 36-2 in dark blue and orange, respectively, and patients not treated with lactulose in grey (n=6 per isolate, results combined from two independent experiments). **D.** Sequence alignment of *lacO* and 5’ end of *lacZ* comparing the reference sequence MP1 with strains 36-1 and 36-2. **E.** Sequence alignment of *lacY* from strains 36-1 and 36-2 was performed with only site of disagreement shown. Data shown as mean ± s.d. Statistical analysis was performed using one-way ANOVA with Bonferroni correction comparing each strain to the control WT *E. coli* strain (**B and C**). *p<0.05, **p<0.01, ***p<0.001, ****p<0.0001.

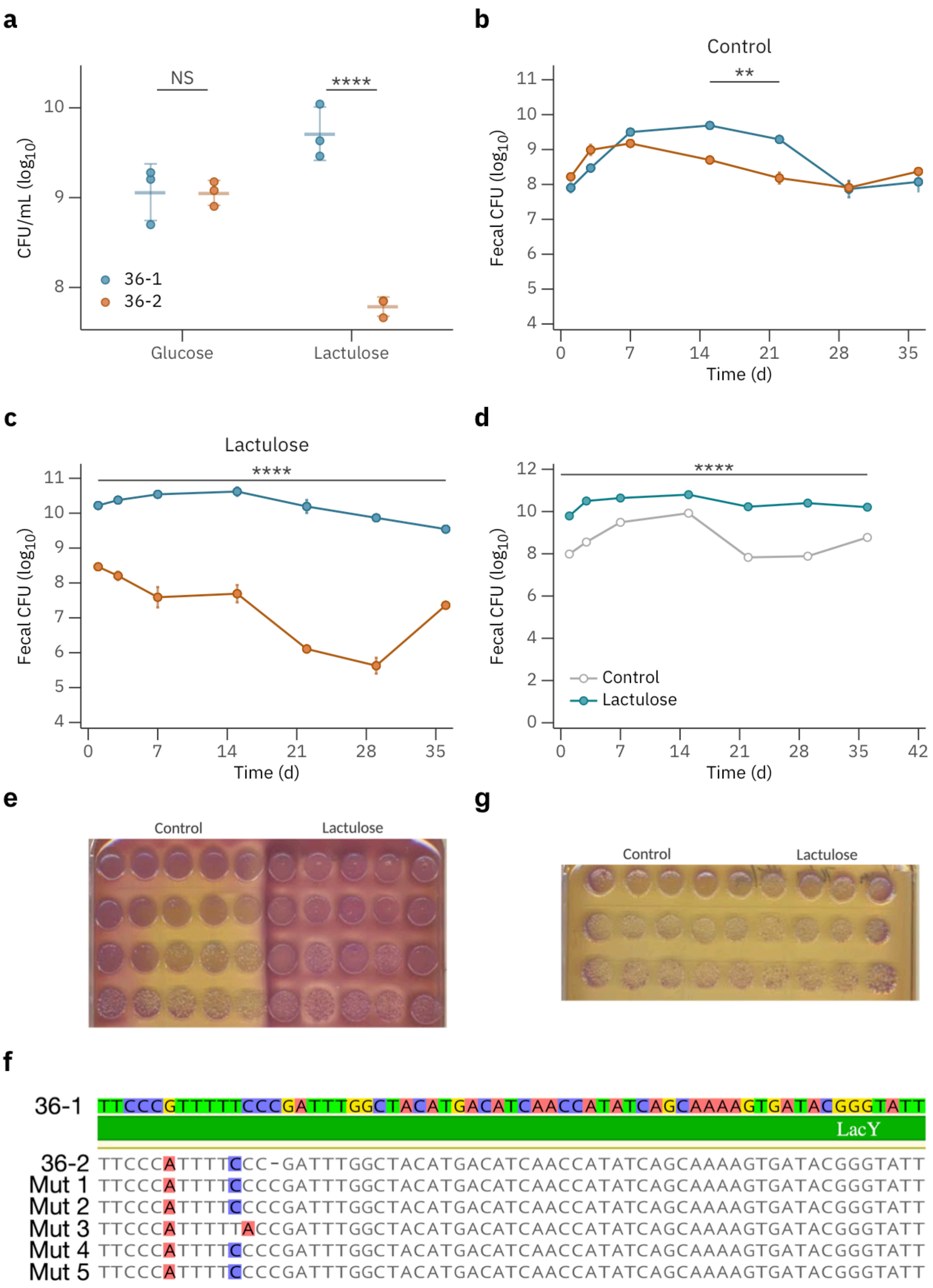
a
NS
****
CFU/mL (log10)
10
9
8
36-1
36-2
Glucose
Lactulose
b
Control
**
Fecal CFU (log10)
11
10
9
8
7
6
5
4
0
7
14
21
28
35
Time (d)
c
Lactulose
****
Fecal CFU (log10)
11
10
9
8
7
6
5
4
0
7
14
21
28
35
Time (d)
d
****
Fecal CFU (log10)
12
10
8
6
4
2
0
Control
Lactulose
0
7
14
21
28
35
42
Time (d)
e
Control
Lactulose
g
Control
Lactulose
f
36-1 TTCCCGTTTTTCCCGATTTGGCTACATGACATCAACCATATCAGCAAAAGTGATACGGGTATT
LacY
36-2 TTCCCATTTTCCC-GATTTGGCTACATGACATCAACCATATCAGCAAAAGTGATACGGGTATT
Mut 1 TTCCCATTTTCCCCGATTTGGCTACATGACATCAACCATATCAGCAAAAGTGATACGGGTATT
Mut 2 TTCCCATTTTCCCCGATTTGGCTACATGACATCAACCATATCAGCAAAAGTGATACGGGTATT
Mut 3 TTCCCATTTTTACCGATTTGGCTACATGACATCAACCATATCAGCAAAAGTGATACGGGTATT
Mut 4 TTCCCATTTTCCCCGATTTGGCTACATGACATCAACCATATCAGCAAAAGTGATACGGGTATT
Mut 5 TTCCCATTTTCCCCGATTTGGCTACATGACATCAACCATATCAGCAAAAGTGATACGGGTATT

**Extended Data Figure 6. A.** *In vitro* competition of strains 36-1 and 36-2, grown in minimal media containing glucose or lactulose (n=3 competitions per condition, results representative of three independent experiments). **B and C.** Fecal CFU of mice colonized with strains 36-1 and 36-2 treated without (**B**) or with (**C**) lactulose (n=5 mice per group, results representative of two independent experiments). **D-F.** Mice were colonized with strain 36-2 and fecal CFU was monitored in the absence or presence of lactulose treatment (**D**). Feces was plated on MacConkey agar with lactulose and colony color was observed (**E**). Individual clones positive for lactulose metabolism underwent WGS and sequence alignment of *lacY* was performed comparing strains 36-1, 36-2, and one representative fecal isolate per mouse (**F**) (n=5 mice per group, results representative of two independent experiments). **G.** Serial dilutions of mouse feces on MacConkey with lactulose selective for strain 36-2 after co-colonization with strains 36-1 and 36-2 in the absence or presence of lactulose treatment (n=5 mice per group, results representative of two independent experiments). Data shown as mean ± s.d. Statistical analysis was performed with ordinary one-way ANOVA with Bonferroni correction for multiple comparisons (**A**) or multiple t-tests with Holm-Sidak correction for multiple comparisons (**B-D**). $^{**}p<0.01$, $^{****}p<0.0001$.

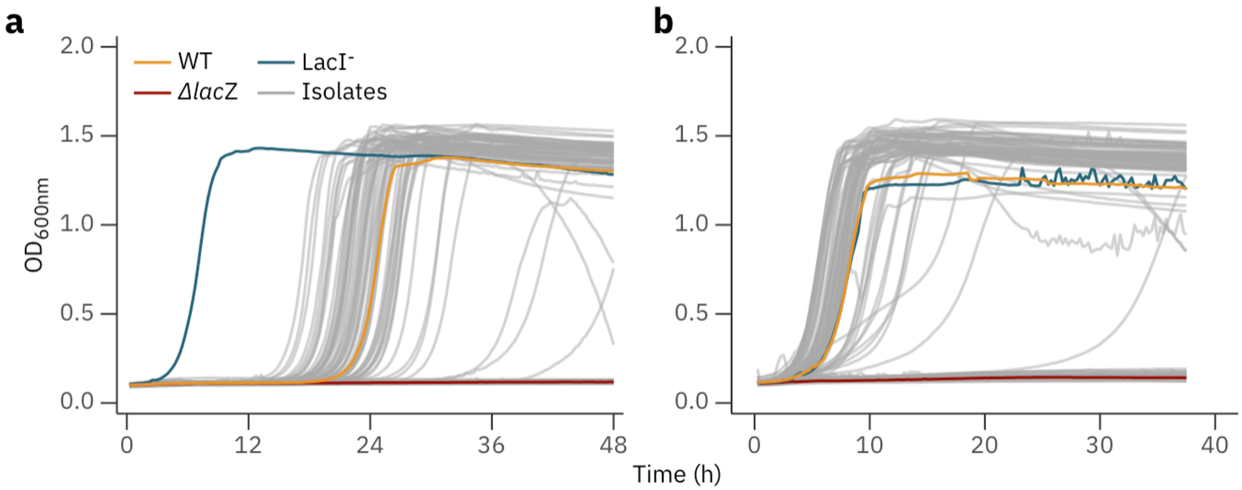


**Extended Data Figure 7. A and B**. Growth curves of *E. coli* bacteremia isolates during first (**A**) and second (**B**) passage of growth in minimal media containing lactulose (n=1 isolate per curve, results representative of two independent experiments). Comparisons made to reference MP13 WT, LacI$^-$ and *ΔlacZ E. coli* strain controls.
